# Multi-layered networks of SalmoNet2 enable strain comparisons of the *Salmonella* genus on a molecular level

**DOI:** 10.1101/2021.12.20.473597

**Authors:** Marton Olbei, Balazs Bohar, David Fazekas, Matthew Madgwick, Padhmanand Sudhakar, Isabelle Hautefort, Aline Métris, Jozsef Baranyi, Robert A. Kingsley, Tamas Korcsmaros

## Abstract

Serovars of the genus *Salmonella* primarily evolved as gastrointestinal pathogens in a wide range of hosts. Some serotypes later evolved further, adopting a more invasive lifestyle in a narrower host range associated with systemic infections. A system-level knowledge of these pathogens has the potential to identify the complex adaptations associated with the evolution of serovars with distinct pathogenicity, host range and risk to human health. This promises to aid the design of interventions and serve as a knowledge base in the *Salmonella* research community. Here we present SalmoNet2, a major update to SalmoNet, the first multi-layered interaction resource for *Salmonella* strains, containing protein-protein, transcriptional regulatory and enzyme enzyme interactions. The new version extends the number of *Salmonella* genomes from 11 to 20, including strains such as *S*. Typhimurium D23580, an epidemic multidrug-resistant strain leading to invasive non-typhoidal *Salmonella* Disease (iNTS), and a strain from *Salmonella bongori*, another species in the *Salmonella* genus. The database now uses strain specific metabolic models instead of a generalised model to highlight differences between strains. This has increased the coverage of high-quality protein-protein interactions, and enhances interoperability with other computational resources by adopting standardised formats. The resource website has been updated with tutorials to help researchers analyse their *Salmonella* data using molecular interaction networks from SalmoNet2. SalmoNet2 is accessible at http://salmonet.org/.

**Importance:** Multi-layered network databases collate information from multiple sources, and are powerful both as a knowledge base and platform for analysis. Here we present SalmoNet2, an integrated network resource of 20 *Salmonella* strains, containing protein-protein, transcriptional regulatory, and metabolic interactions. Key improvements to the update include expanding the number of strains, strain-specific metabolic networks, an increase in high quality protein-protein interactions, community standard computational formats to help interoperability, and online tutorials to help users analyse their data using SalmoNet2.

## Introduction

Serovars of the genus *Salmonella* are enteric pathogens, capable of causing a self-limiting gastrointestinal inflammatory disease in a variety of animals. The host species, depending on the *Salmonella* subspecies, range from cold-blooded vertebrates to humans. *Salmonella* infection is one of the most common foodborne or waterborne illnesses resulting in approximately 94 million illnesses, and 155,000 deaths each year ^1–3^.

Of six subspecies of *Salmonella enterica*, a small number of subspecies I serovars have adapted to cause an invasive infection in a restricted host range, instead of a self-limiting gastrointestinal inflammation typical of *Salmonella* serovars. These extraintestinal *Salmonella* strains, including the typhoidal strains that are human adapted, emerged on multiple occasions independently. A hallmark of adaptation is genomic and phenotypic changes, including loss of function mutations in genes related to adaptation to specific niches in their host commonly affecting anaerobic metabolism, virulence genes, chemotaxis or motility ^4^

*S*. Typhi is an ancient pathogen and the most common extraintestinal *Salmonella* serovar affecting humans. Over the past decades, another group of *Salmonella* appeared as one of the most commonly isolated pathogens from the blood of patients, particularly in sub-Saharan Africa ^5^. The invasive nontyphoidal *Salmonella* (iNTS) strains, in common with *S.* Typhi and *S*. Paratyphi, cause a systemic infection. Unlike *S*. Typhi, iNTS commonly affects immunocompromised individuals or young children, leading to bacteremia and meningitis. iNTS is most often caused by specific genotypes of *S*. Typhimurium and *S*. Enteritidis that are distinct from genotypes of these serovars commonly associated with gastrointestinal infections outside of sub-Saharan Africa ^6–8^.

The *Salmonella* genus contains pathogens with diverse host range and pathogenicity, and dissecting the specific differences between gastrointestinal and extraintestinal strains have been pursued by a multitude of means ^4,9,10^ Previously, we constructed SalmoNet, a multi-layered network resource for 10 *Salmonella* serovars that integrated protein-protein, regulatory and metabolic information ^11^. With its multi-layered networks SalmoNet can serve as a knowledge base for the community and aid in understanding *Salmonella* pathogenesis and evolution by mapping the differences in molecular interactions between *Salmonella* pathovars on multiple biological levels. This systems level information allows researchers to enhance the information content of their own studies, by adding interaction context to the changes observed on a genomic or transcriptome level.

Here we present SalmoNet2, an update to the first public multi-layered network resource for *Salmonella* research. The new version extends the coverage of strains from 11 to 20, including an important iNTS strain, and strains outside of subspecies *enterica*, from subspecies *arizonae* and *Salmonella bongori*. To aid interoperability in computational biology the database adopted the PSI-MI TAB format, and is now accessible through the NDEx network repository. In addition, we show how rewiring of the network information can be utilised by the research community to understand aspects of *Salmonella* evolution, the step-by-step workflows of which are now accessible through tutorials on our website.

## Results

### SalmoNet2 extends out of subspecies I

SalmoNet2 adds 9 new multi-layered networks of *Salmonella* strains in the database compared to the first version. Included amongst others are commonly used laboratory strains, additional extraintestinal strains, including *S*. Typhimurium strain D23580, a well characterised pathogen associated with invasive non-typhoidal *Salmonella* (iNTS) disease, a strain of *Salmonella bongori*, and a member of a different subspecies (subsp. *arizonae*) within *Salmonella enterica*. The extended coverage captures a larger variety of the *Salmonella* genus, and for the first time provides interaction networks for strains from outside of subspecies *enterica* (Supplementary Table I). To define the phylogenetic relationship of the strains included in the database we constructed a neighbour-joining tree from variation in the core genome nucleotide sequence, and compared this with hierarchical classification trees based on matrix representation of protein-protein, regulatory and metabolic networks (Figure 1).

**Figure 1.**
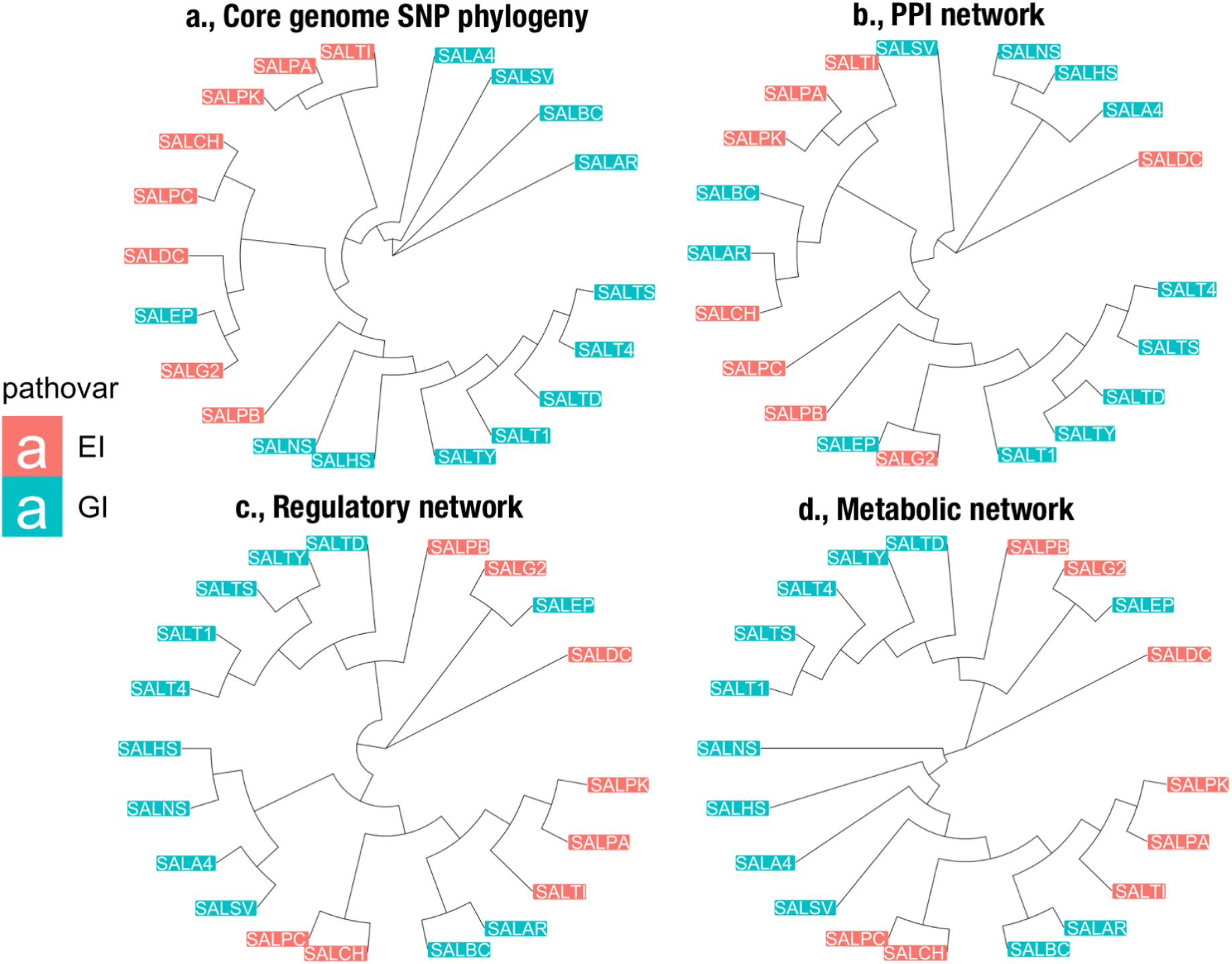
Core genome SNP based phylogenetic tree, and hierarchical classification of network layers. Extraintestinal (EI) serovars labelled with red, gastrointestinal (GI) serovars with blue labels. A., Neighbour-joining tree from core genome SNPs of the strains. B-D., Hierarchical classification trees based on matrix representation of protein-protein, regulatory and metabolic networks. The five letter labels encode for the names of the different strains (for details of the encoding please refer to Supplementary Table 1).

The topology was in accordance with previously published phylogenies^12^ with no clear clustering of extraintestinal and gastrointestinal serovars in the phylogenetic tree. This is consistent with observations in previous works in the literature, where the extraintestinal and gastrointestinal strains could not be distinguished based on genomic dendrograms, and consistent with the independent emergence of extraintestinal serovars from gastrointestinal serovars, through a convergent evolutionary process ^13,14^

### SalmoNet2 increases the information content of the individual network layers

We included a number of methodological improvements to the workflow of the Salmonet1 database, leading to an increased number of high quality interactions for the individual network layers. To increase the coverage of the protein-protein interactions without compromising quality, we have used the IntAct MIscore when extrapolating orthologous interaction information from the IntAct database ^15^. Instead of relying on one experimental method as in the first version, using the MIscore as a quality filter permited extending the number of available high-quality protein-protein interactions that we could use to establish orthologous protein-protein interactions from the commensal bacteria *Escherichia coli* (Supplementary Figure 1).

By utilising a strain-specific genome-scale metabolic model for each strain developed previously (*Seif et al*.), instead of a general model (*Thiele et al*.), the metabolic layer now includes more enzyme-enzyme relationships, where two proteins are connected if a metabolite produced by one is a substrate for another ^16–18^, leading to a more complete description of the metabolic capabilities of the strains. The information content of Position-Specific Scoring Matrices (PSSMs) that are required to carry out genome-wide regulatory scans were enhanced with novel binding sites published since the first version of the database, and from new data uploaded to the CollecTF repository ^19^. The total number of interactions has been increased from 81,514 to 270,215, primarily due to the expansion of the PPI layer, and the increase in the number of involved strains. The composition of the consensus network, comprised of shared interactions amongst all strains included in the database, slightly changed from the first version of SalmoNet, indicating the shifts caused by the updated data sources and expanded strain repertoire. 24.4% of regulatory interactions (up from 16%), 68.1% of PPI interactions (down from 72%), and 51.8% (down from 69%) of metabolic interactions were shared amongst all strains, forming the core network of *Salmonella* interactions. Figure 2 shows the changes in the size of the networks and individual layers compared to the first version.

**Figure 2.**
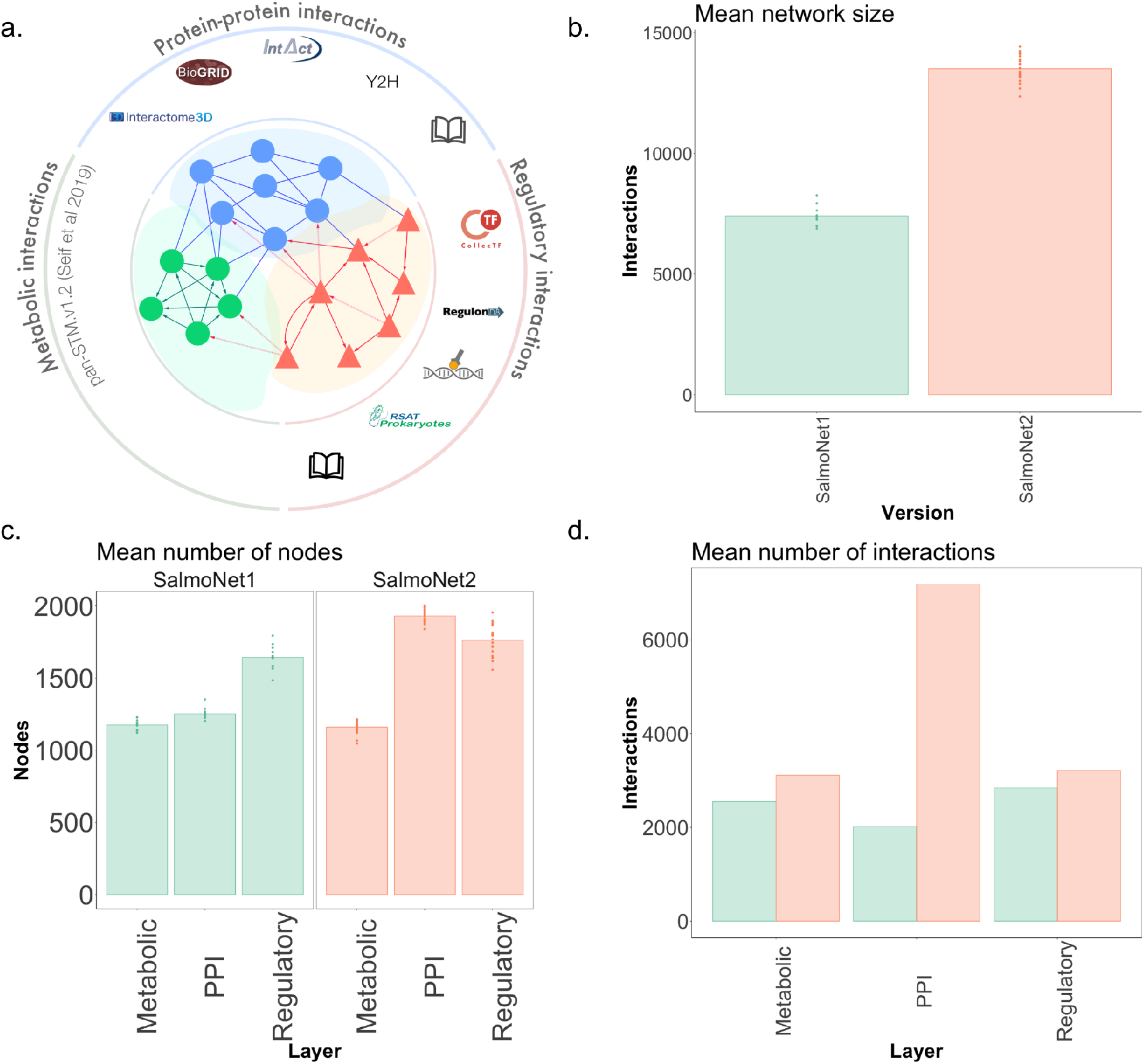
Comparison of SalmoNet2 with the first version. A: main data sources and interactions in SalmoNet2. B: comparison of network size in SalmoNet 1 and SalmoNet2. C: comparison of layer size in terms of participating nodes. D: comparison of layer size in terms of interactions between SalmoNet and SalmoNet2. The five letter codes encoding for the different strains can be found in Supplementary Table 1.

### Novel formats improve interoperability

In addition to the previously used formats (.csv, .cys.), we extended the output format data to help computational biologists access network information in SalmoNet2. We now provide networks in the community standard PSI-MITAB format as well, which contains a strictly regulated vocabulary for interaction data, helping interoperability between network resources, a prerequisite for inclusion in the PSICQUIC ecosystem ^20^. Using standardised formats improves the interoperability with other network information repositories, and provides space to maximise each interaction with as much data as possible, in a controlled and transparent manner ^20^. This further strengthened the information content of the database, and improved the potential use cases beyond network analysis. To enable the networks to be directly accessible from the widely used Cytoscape network analysis program, we have also deposited them to the NDEx network repository ^21^.

### Website enhancements for a user-friendly experience

The SalmoNet website was enhanced compared to the previous version. We now carry new locus tag identifiers for all *Salmonella* strains to enable users to map their experimental data to the SalmoNet2 interaction networks. As part of our shift to OMA as the source of orthology for *Salmonella* proteins, SalmoNet2 now directly links to the respective OMA pages and sequence data instead of Uniprot ^22^. Where possible, Uniprot data is still accessible through OMA ^23^.

During the lifecycle of SalmoNet1, we identified a bottleneck with our putative users. The interaction network format, while potentially useful for scientists with a microbiology background, proved difficult to use, which led to potentially less user retention. To combat this, we have created a new tab on the website containing tutorials as an introduction to network analysis using the SalmoNet2 database. These tutorials enable analyses shown in this article. We plan to add additional tutorials, workflows and examples to the website in the future, to further increase the usability of the platform.

### Case study: network rewiring to identify functional differences in *Salmonella enterica*

Network rewiring entails many approaches aimed at quantifying changes between interaction networks, and has been used to identify differences between interaction networks ^24,25^. In this work, we compared the degree of interaction rewiring between the interactomes of four host adapted typhoidal *Salmonella* strains and four gastrointestinal *Salmonella* strains to explore the utility of a multi-layered network resource such as SalmoNet2. We compared the most rewired subgraphs of the two types of strains to find the causes of the rewiring.

In general, the most rewired nodes were global regulators, such as Crp, Fis and Fur. The significantly enriched functions are similar between the compared strains, with a few key differences. For example, the ferric uptake regulator Fur senses metal concentration and redox state of cells, and regulates many operons and genes involved in these processes ^26^. Interestingly, Fur is enriched in the GO function “iron ion homeostasis” in all included gastrointestinal strains, while this enrichment is absent from the typhoidal strains. Upon further inspection of the genes responsible for the enrichment of the term and their orthologous status, Fur is missing interactions present in GI strains towards the genes *fhuA, fhuE,* caused by the disruption of coding sequences in these genes in typhoidal serovars, as highlighted previously in the literature ^13,27^ Similarly, Fur is enriched in the term “cell adhesion” in all gastrointestinal strains, whereas this function is not enriched in typhoidal strains, except *S*. Paratyphi C. Inspection of the genes underlying the enrichment result revealed that the culprit behind the mismatch in functional enrichment is the pseudogenization and subsequent missing interactions with the genes *stiH* and *stiA* in the rest of the typhoidal *Salmonella* strains, two genes responsible for the production of fimbriae, highlighted previously in the literature ^13^. From the top 50 most rewired nodes, on average 33 nodes had at least one pseudogene first neighbour in the typhoidal serovars, and on average 4% of the first neighbours of the top 50 most rewired nodes were pseudogenes. In the gastrointestinal strains, on average 7 nodes had pseudogene first neighbours, and only 1% of their first neighbours were pseudogenes.

While a large part of the rewiring was due to gene loss in typhoidal and extraintestinal serovars, we found examples where the cause of rewiring was due to the exclusivity of genes to the extraintestinal group. Two proteins, YreP and YjcS, are present in all typhoidal and extraintestinal strains of *Salmonella* included in SalmoNet2. However, they are missing from all gastrointestinal strains bar one. The protein YjcS has an orthologue in *S*. Enteritidis, but the protein is otherwise missing from the gastrointestinal group. The genes share an upstream regulatory region, and are predicted to interact with the regulators HilC, RtsA and Fur. The *yreP* and *yjcS* genes were first described together in *Escherichia coli*, in two analysed strains: *E. coli* SMS-3-5, an environmental pathogenic isolate with multiple antibiotic resistances, and *E. coli* (NMEC) O7:K1 strain CE10, causing neonatal meningitis. The first gene, *yreP* (*dgcY* in *E. coli*), encodes a putative diguanylate cyclase, based on the presence of a GGDEF domain ^28,29^. Diguanylate-cyclases facilitate the production of c-di-GMP, a ubiquitous secondary messenger metabolite in prokaryotes ^28,29^. The second gene, *yjcS* (EcSMS35_1714 in *E. coli*), is an alkyl-sulfatase. This enzyme has been first described in *Pseudomonas spp*., where a strain carrying this enzyme was able to grow on the surfactant sodium dodecyl sulphate (SDS), and the gene has been characterised in *E. coli* as well ^30,31^.

After noting their presence in the extraintestinal strains included in SalmoNet2, we expanded the search into a more expansive data source, pubMLST, to see if this split was representative of the serovars as a whole, and not just the specific strains in SalmoNet2 ^32^. Figure 3 shows the results of the BLAST searches in the pubMLST database.

**Figure 3.**
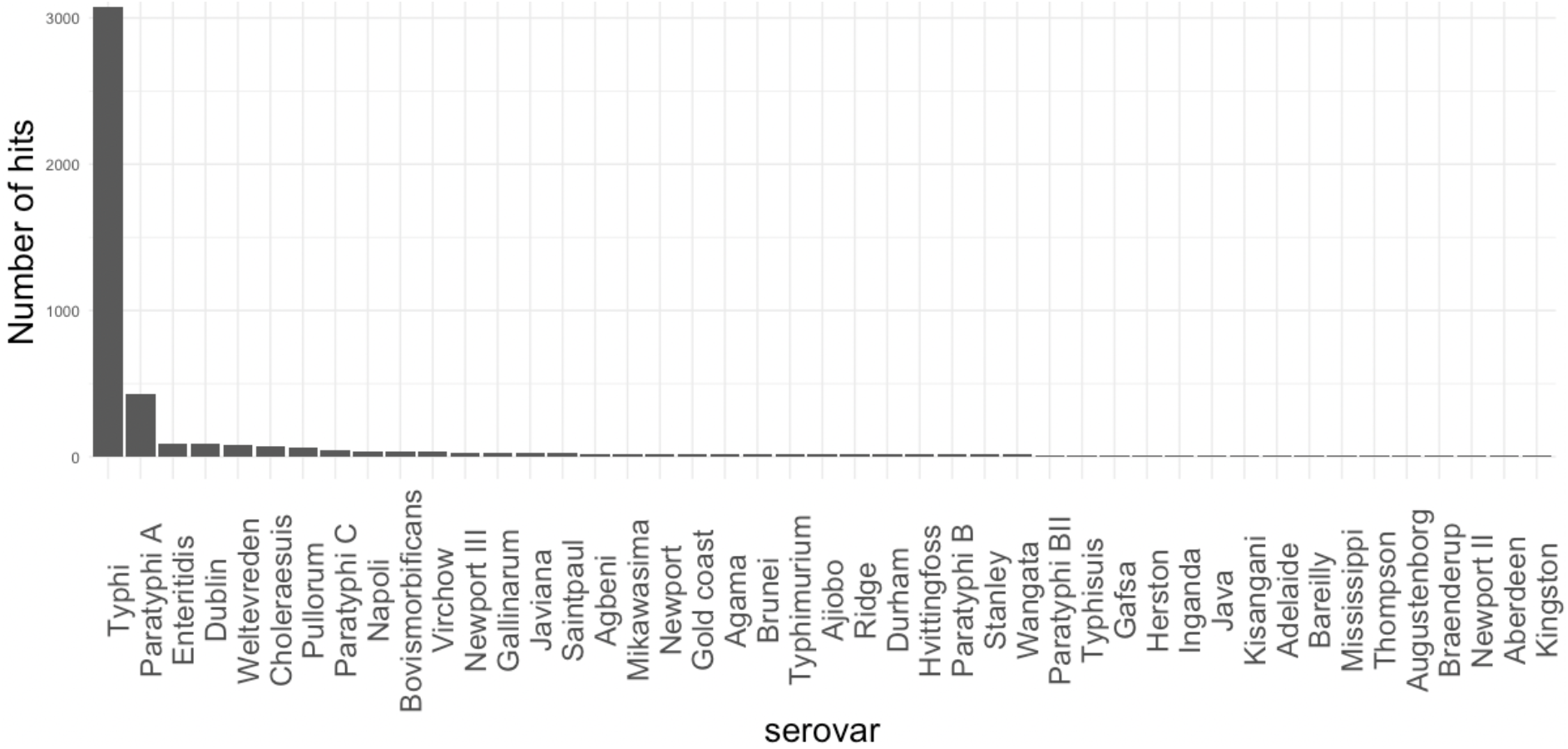
Prevalence of the yreP + regulatory region + yjcS segment in Salmonella serovars based on BLAST hits. The top 10 serovars have been described previously as sources of invasive illness. Serovars containing < 5 isolates were removed from this figure for clarity.

In total 83% of BLAST hits come from well-known extraintestinal serovars, dominated by *S*. Typhi strains (Figure 3). The top 10 serovars in terms of number of hits are mostly invasive serovars: *S*. Typhi, *S*. Paratyphi A, and *S*. Paratyphi C are notable typhoidal serovars adapted to humans, *S*. Dublin, *S*. Pullorum and *S*. Choleraesuis are well-known host adapted serovars of cattle, poultry and pigs ^4,11^. *S*. Napoli is an emerging serovar in Europe, phylogenetically closely related to *S*. Paratyphi A, carrying an almost identical pattern of typhoid-associated genes, and capable of causing a form of invasive non-typhoidal disease ^33,34^. The invasive behaviour is not as clear cut with the rest of the serovars, but there have been several reports of it: *S*. Bovismorbificans is capable of causing bloodstream infections, and has recently been described as an emerging disease in Malawi, converging towards a phenotype resembling a human adapted iNTS variant ^35^. Although not strictly an extraintestinal serovar, *S*. Virchow has been known to cause invasive illness ^36–39^. *S*. Weltevreden is an emerging cause of diarrheal and sometimes invasive disease in humans in tropical regions, and may be adapted to life in aquatic hosts ^40,41^. While large in total numbers in the database, *S*. Enteritidis only makes up 2% of the positive hits. Since *S*. Enteritidis is one of the most commonly isolated iNTS strains, there exists a possible link to invasive behaviour ^42,43^. However, more work is needed to uncover whether the two proteins are beneficial to an extraintestinal lifestyle.

This brief case study highlights how the information contained in and linked with SalmoNet2 can be used to form scientific questions relating the functionality of genes to the behaviour and phylogenetics of *Salmonella*, based on molecular interaction information. SalmoNet 2 contains example strains from the most prevalent serovars, and the information can further be extended using the easily accessible sequence data and homology information through OMA and other computational resources.

## Discussion

By increasing the number of strains to 20 from the previous 11, SalmoNet now extends out of subspecies I., adding information on members of other subspecies (subspecies *arizonae*), or an entirely different species (*Salmonella bongori*). Developing a more compatible structure between SalmoNet, and other available large-scale evolutionary genomics tools such as OMA, there is increased potential to generate interaction networks for specific *Salmonella* strains on request, or build similar data resources for other non-model organisms ^44^. With the change to OMA as the backbone of SalmoNet interactions, there is a great potential to study the evolutionary history of proteins, and interactions. The on-demand availability of orthologous proteins from outside of the studied organism or clade can make larger scope comparisons possible ^45^.

The programmatic access interfaces implemented into OMA make these integrated analyses reproducible and scalable^46^. Orthology mapping is the most computationally intensive step of the SalmoNet workflow. The OMA standalone software can save a lot of time and resources here, since the all-against-all Smith-Waterman sequence alignments can be parallelised, both on single computers or high-performance clusters ^22^. Adding a new strain or species in the future is also made easier, as OMA Standalone does not require an all-against-all recomputing of the orthologous relationships in these cases, as pre-computed results can be submitted, in which case only the new genomes require computation time. Using OMA is not only beneficial for the orthology mapping, it is also helpful for the annotation work. The first version of SalmoNet was essentially UniProt based, with UniProt IDs serving as the primary identifiers of the database. Currently not all proteins of all strains have a matching UniProt ID, hence the OMA IDs as our new primary identifier.

The availability of strain-specific metabolic models, and the increased specificity of PPI data, although still reliant on orthology mapping, increases the resolution of the resulting network models for non-model organisms, and the more interwoven interaction layers get, the more valuable the information content of the database gets. Although there are other resources containing *Salmonella* interaction data, such as STRING for PPI interactions, RegPrecise for regulatory interactions, or BioCyc for metabolic interactions, no other freely available resource combines the listed connection types besides SalmoNet, for multiple *Salmonella* strains ^47–49^.

SalmoNet2 enables the network analyses as shown with the rewiring analysis. It highlights how the information contained in and linked with SalmoNet2 can be used to inform scientific questions such as relating the functionality of genes to phenotypes and phylogenetics of *Salmonella*, based on molecular interaction information. SalmoNet2 contains example strains from the most prevalent serovars, and the information can further be extended using the easily accessible sequence data and homology information through OMA and other computational resources as shown with the pubMLSt example.

To increase the usability and interoperability of the generated interaction information, we now utilise the PSI-MITAB format as well, quickly becoming a standard of biological network information ^20,50^. To have the networks be directly accessible from the widely used Cytoscape network analysis program, the NDEx network repository can host them separately from the SalmoNet website, making them directly available for end-users ^21^. Beyond their raw information content, databases are as good as their usability and availability, and the potential for SalmoNet2 data to be found and utilised in as many ways as possible is crucial for this effort to be useful for the scientific community ^51^. To further enhance the accessibility of SalmoNet2 data we wrote detailed step-by-step tutorials describing the computational steps required to perform analyses such as the comparisons involving the gastrointestinal and typhoidal strains above.

## Methods

### Updated orthology mapping tool

Although the main structure of the database remained the same, the underlying workflow changed. As in the first version, we mapped the orthologous proteins across the included strains. In SalmoNet 1 this was done by InParanoid, a well-established tool for this process ^52^. In this update we used the OMA standalone software to construct these relationships, including the available *Salmonella* strains from the OMA browser database. OMA is a large-scale orthology database and toolkit, containing the orthology information and protein sequence data needed for SalmoNet in one place, including the proteomes and genomes of the strains on request, and important annotation data ^53^.

It is important to note, that the outputs of the tools can be slightly different: according to a study comparing these methods the OMA standalone output OMA groups lead to a generally more precise, but also strict mapping, leading to less false positives (and true positives as well) ^54^. We did however get very similar, and in cases better recall than we did in SalmoNet 1.0 (between 69-75% overlap with the 4140 proteins from *E. coli*; Supplementary Table 1) using InParanoid.

### Updated and novel data sources

#### Protein-Protein Interaction Networks

The construction of the protein-protein interaction (PPI) network follows the same essential steps it did in the first version of the database, collected from multiple databases ^55–58^. To increase the coverage of the included PPIs without losing quality, we have used the IntAct PSI-MIscore (> 0.50) when importing interaction information from the IntAct database, instead of relying on one experimental method, as in the first version (psi-mi:“MI:0096”(pull down)). Supplementary Figure 1 shows the distribution of the IntAct PSI-MIscores.

#### Metabolic Networks

SalmoNet2 uses new, strain specific genome-scale metabolic models for *Salmonella* ^16,17^ The models used the same STM 1.0 model as a starting point SalmoNet1 did ^18^, but updated it with new genes and reactions, and were made strain specific, leading to the metabolic models of 410 *Salmonella* strains belonging to 64 serovars. Otherwise, the workflow remained identical, resulting in enzyme-enzyme interactions, where two proteins are connected if a metabolite produced by one is a substrate for another ^59^. Similarly, as in the first version, we have excluded links connected by metabolites partaking in more than 10 reactions ^59^.

#### Regulatory Networks

The establishment of the transcriptional regulatory networks was done in an identical way to SalmoNet 1. Supplementary Figure 2 shows the workflow for the construction of the regulatory layer. The core of the network, the manually curated interactions, high-throughput data (ChIP-Seq), and low-throughput, experimentally verified interactions and data sources remained the same. The information content of Position-Specific Scoring Matrices (PSSMs) used to carry out the genome-wide scans was enhanced with novel binding sites published since the first version of the database, from new data uploaded to the CollecTF repository ^19^. RSAT’s consensus tool is no longer available on the web server, info-gibbs took its place, which is the tool that was used to construct the matrices. Similarly, as previously, RSAT retrieve-sequence was used to gather the putative promoter regions for the genomes included in SalmoNet, and matrix-scan was used to establish putative transcription factor - target gene (promoter region) pairs ^60^.

### Removal of pseudogenes

To remove all hypothetically disrupted coding DNA sequences (HDCs), the curation made by Nuccio & Bäumler was used to remove such entries ^13^ and ^61,62^ were used to remove them from S. Typhimurium D23580.

### Network rewiring

To calculate network rewiring we used the DyNet app in Cytoscape to calculate the rewiring value of the nodes in each group separately ^63^. Four typhoidal strains (S. Paratyphi A (AKU 1261), S. Paratyphi A (ATCC 9150), S. Paratyphi C (RKS4594), S. Typhi (Ty2) and four gastrointestinal strains (S. Agona (SL483), S. Newport (SL254), S. Heidelberg and S. Typhimurium (LT2)) were compared for interaction differences.

The level of rewiring was calculated across all strains, and the degree-corrected rewiring values were ordered in a descending list, where the top 50 hits were further analysed. To calculate the enrichment of Gene Ontology terms in the identified subgraphs the the up-to-date Gene Ontology annotation of the target genes was downloaded using the topGO library in R, and following that the R library clusterProfiler was used to calculate Gene Ontology enrichment with the enricher function, from Biological Process terms ^64,65^. P-value adjustment for multiple testing was carried out with the Benjamini-Hochberg approach, using the p.adjust function in R.

The statistically significant enrichment results were compared side-by-side between the groups, and the differences in enrichment were further studied by comparing the sets of genes responsible for (underlying) the enriched terms, i.e., if one group was enriched in a specific term, the presence/absence of the orthologous genes responsible for the enrichment was analysed in the members of the other group.

To study the relationship of YreP and YjcS to the extraintestinal pathovar, network rewiring was calculated in an identical manner as above, but all extraintestinal and gastrointestinal strains from SalmoNet2 were involved in the comparisons. BLAST searches for the *yreP* and *yjcS* genes was done through the pubMLST website, with default parameters ^32^. The entire genomic sequence of the genes and their shared regulatory region was queried, as taken from *S.* Gallinarum strain 287/91 (see Supplementary File 1). The hits were filtered for above 95% sequence identity, and the top 10% of bitscores to make sure the compared sequences contain both the genes and the shared regulatory region.

## Data availability

The data generated for this study is available at the database website, http://salmonet.org.

## Acknowledgements

The work of MO, PS, IH, TK were supported by the UKRI BBSRC Gut Microbes and Health Institute Strategic Programme BB/R012490/1 and its constituent projects BBS/E/F/000PR10353 and BBS/E/ F/000PR10355. MO, BB, DF, PS, IH, TK were also supported by a BBSRC Core Strategic Programme Grant for Genomes to Food Security (BB/CSP1720/1) and its constituent work packages, BBS/E/T/000PR9819 and BBS/E/T/000PR9817. PS was supported by the European Research Council Advanced Grant (ERC-2015-AdG, 694679, CrUCCial). MO and MM were supported by a BBSRC - Norwich Research Park Biosciences Doctoral Training Partnership grant (BB/M011216/1 and BB/S50743X/1). RK was supported by the UKRI Institute Strategic Programme Microbes in the Food Chain BB/R012504/1 and its constituent project(s) BBS/E/F/000PR10348 and BBS/E/F/000PR10349.

## Bibliography

1. Coburn, B., Grassl, G. A. & Finlay, B. B. Salmonella, the host and disease: a brief review. Immunol. Cell Biol. 85, 112–118 (2007).

2. Hohmann, E. L. Nontyphoidal salmonellosis. Clin. Infect. Dis. 32, 263–269 (2001).

3. Majowicz, S. E. et al. The global burden of nontyphoidal Salmonella gastroenteritis. Clin. Infect. Dis. 50, 882–889 (2010).

4. Tanner, J. R. & Kingsley, R. A. Evolution of Salmonella within Hosts. Trends Microbiol. 26, 986–998 (2018).

5. Tsai, C. N. & Coombes, B. K. Emergence of invasive Salmonella in Africa. Nat. Microbiol. (2021) doi:10.1038/s41564-021-00864-5.

6. Gilchrist, J. J. & MacLennan, C. A. Invasive nontyphoidal salmonella disease in africa. Ecosal Plus 8, (2019).

7. Feasey, N. A., Dougan, G., Kingsley, R. A., Heyderman, R. S. & Gordon, M. A. Invasive non-typhoidal salmonella disease: an emerging and neglected tropical disease in Africa. Lancet 379, 2489–2499 (2012).

8. Okoro, C. K. et al. Intracontinental spread of human invasive Salmonella Typhimurium pathovariants in sub-Saharan Africa. Nat. Genet. 44, 1215–1221 (2012).

9. Perez-Sepulveda, B. M. & Hinton, J. C. D. Functional transcriptomics for bacterial gene detectives. Microbiol. Spectr. 6, (2018).

10. Langridge, G. C., Nair, S. & Wain, J. Nontyphoidal*salmonella* serovars cause different degrees of invasive disease globally. J. Infect. Dis. 199, 602–603 (2009).

11. Métris, A. et al. SalmoNet, an integrated network of ten Salmonella enterica strains reveals common and distinct pathways to host adaptation. NPJ Syst. Biol. Appl. 3, 31 (2017).

12. Branchu, P., Bawn, M. & Kingsley, R. A. Genome Variation and Molecular Epidemiology of Salmonella enterica Serovar Typhimurium Pathovariants. Infect. Immun. 86, (2018).

13. Nuccio, S.-P. & Bäumler, A. J. Comparative analysis of Salmonella genomes identifies a metabolic network for escalating growth in the inflamed gut. MBio 5, e00929–14 (2014).

14. Timme, R. E. et al. Phylogenetic diversity of the enteric pathogen Salmonella enterica subsp. enterica inferred from genome-wide reference-free SNP characters. Genome Biol. Evol. 5, 2109–2123 (2013).

15. Villaveces, J. M. et al. Merging and scoring molecular interactions utilising existing community standards: tools, use-cases and a case study. Database (Oxford) 2015, (2015).

16. Seif, Y., Monk, J. M., Machado, H., Kavvas, E. & Palsson, B. O. Systems Biology and Pangenome of Salmonella O-Antigens. MBio 10, (2019).

17. Seif, Y. et al. Genome-scale metabolic reconstructions of multiple Salmonella strains reveal serovar-specific metabolic traits. Nat. Commun. 9, 3771 (2018).

18. Thiele, I. et al. A community effort towards a knowledge-base and mathematical model of the human pathogen Salmonella Typhimurium LT2. BMC Syst. Biol. 5, 8 (2011).

19. Kiliç, S. et al. From data repositories to submission portals: rethinking the role of domain-specific databases in CollecTF. Database (Oxford) 2016, (2016).

20. Perfetto, L. et al. CausalTAB: the PSI-MITAB 2.8 updated format for signalling data representation and dissemination. Bioinformatics 35, 3779–3785 (2019).

21. Pillich, R. T., Chen, J., Rynkov, V., Welker, D. & Pratt, D. Ndex: A community resource for sharing and publishing of biological networks. Methods Mol. Biol. 1558, 271–301 (2017).

22. Altenhoff, A. M. et al. The OMA orthology database in 2018: retrieving evolutionary relationships among all domains of life through richer web and programmatic interfaces. Nucleic Acids Res. 46, D477–D485 (2018).

23. The UniProt Consortium. UniProt: the universal protein knowledgebase. Nucleic Acids Res. 45, D158–D169 (2017).

24. Mehta, T. K. et al. Evolution of regulatory networks associated with traits under selection in cichlids. Genome Biol. 22, 25 (2021).

25. Treveil, A. et al. Regulatory network analysis of Paneth cell and goblet cell enriched gut organoids using transcriptomics approaches. Mol. Omics 16, 39–58 (2020).

26. Troxell, B., Fink, R. C., Porwollik, S., McClelland, M. & Hassan, H. M. The Fur regulon in anaerobically grown Salmonella enterica sv. Typhimurium: identification of new Fur targets. BMC Microbiol. 11, 236 (2011).

27. Wang, Y. et al. Evolution and sequence diversity of fhua in salmonella and escherichia. Infect. Immun. 86, (2018).

28. Povolotsky, T. L. & Hengge, R. Genome-Based Comparison of Cyclic Di-GMP Signaling in Pathogenic and Commensal Escherichia coli Strains. J. Bacteriol. 198, 111–126 (2016).

29. Ryjenkov, D. A., Tarutina, M., Moskvin, O. V. & Gomelsky, M. Cyclic diguanylate is a ubiquitous signaling molecule in bacteria: insights into biochemistry of the GGDEF protein domain. J. Bacteriol. 187, 1792–1798 (2005).

30. Liang, Y., Gao, Z., Dong, Y. & Liu, Q. Structural and functional analysis show that the Escherichia coli uncharacterized protein YjcS is likely an alkylsulfatase. Protein Sci. 23, 1442–1450 (2014).

31. Williams, J. & Payne, W. J. Enzymes induced in a bacterium by growth on sodium dodecyl sulfate. Appl. Microbiol. 12, 360–362 (1964).

32. Jolley, K. A., Bray, J. E. & Maiden, M. C. J. Open-access bacterial population genomics: BIGSdb software, the PubMLST.org website and their applications. [version 1; peer review: 2 approved]. Wellcome Open Res. 3, 124 (2018).

33. Gori, M. et al. High-resolution diffusion pattern of human infections by Salmonella enterica serovar Napoli in Northern Italy explained through phylogeography. PLoS ONE 13, e0202573 (2018).

34. Huedo, P. et al. Salmonella enterica Serotype Napoli is the First Cause of Invasive Nontyphoidal Salmonellosis in Lombardy, Italy (2010-2014), and Belongs to Typhi Subclade. Foodborne Pathog. Dis. 14, 148–151 (2017).

35. Bronowski, C. et al. Genomic characterisation of invasive non-typhoidal Salmonella enterica Subspecies enterica Serovar Bovismorbificans isolates from Malawi. PLoS Negl. Trop. Dis. 7, e2557 (2013).

36. Eckerle, I., Zimmermann, S., Kapaun, A. & Junghanss, T. Salmonella enterica serovar Virchow bacteremia presenting as typhoid-like illness in an immunocompetent patient. J. Clin. Microbiol. 48, 2643–2644 (2010).

37. Mani, V., Brennand, J. & Mandal, B. K. Invasive illness with Salmonella virchow infection. Br. Med. J. 2, 143–144 (1974).

38. Messer, R. D., Warnock, T. H., Heazlewood, R. J. & Hanna, J. N. Salmonella meningitis in children in far north Queensland. J. Paediatr. Child Health 33, 535–538 (1997).

39. Todd, W. T. & Murdoch, J. M. Salmonella virchow: a cause of significant bloodstream invasion. Scott. Med. J. 28, 176–178 (1983).

40. Hounmanou, Y. M. G. et al. Molecular characteristics and zoonotic potential of salmonella weltevreden from cultured shrimp and tilapia in vietnam and china. Front. Microbiol. 11, 1985 (2020).

41. Makendi, C. et al. A Phylogenetic and Phenotypic Analysis of Salmonella enterica Serovar Weltevreden, an Emerging Agent of Diarrheal Disease in Tropical Regions. PLoS Negl. Trop. Dis. 10, e0004446 (2016).

42. Feasey, N. A. et al. Distinct Salmonella Enteritidis lineages associated with enterocolitis in high-income settings and invasive disease in low-income settings. Nat. Genet. 48, 1211–1217 (2016).

43. Gordon, M. A. Invasive nontyphoidal Salmonella disease: epidemiology, pathogenesis and diagnosis. Curr. Opin. Infect. Dis. 24, 484–489 (2011).

44. Olbei, M., Kingsley, R. A., Korcsmaros, T. & Sudhakar, P. Network Biology Approaches to Identify Molecular and Systems-Level Differences Between Salmonella Pathovars. Methods Mol. Biol. 1918, 265–273 (2019).

45. Demeter, A. et al. ULK1 and ULK2 are less redundant than previously thought: computational analysis uncovers distinct regulation and functions of these autophagy induction proteins. Sci. Rep. 10, 10940 (2020).

46. Kaleb, K., Warwick Vesztrocy, A., Altenhoff, A. & Dessimoz, C. Expanding the Orthologous Matrix (OMA) programmatic interfaces: REST API and the *OmaDB* packages for R and Python [version 2; peer review: 2 approved]. F1000Res. 8, 42 (2019).

47. Caspi, R. et al. BioCyc: A Genomic and Metabolic Web Portal with Multiple Omics Analytical Tools. The FASEB Journal (2019).

48. Novichkov, P. S.et al. RegPrecise 3.0--a resource for genome-scale exploration of transcriptional regulation in bacteria. BMC Genomics 14, 745 (2013).

49. Szklarczyk, D. et al. STRING v11: protein-protein association networks with increased coverage, supporting functional discovery in genome-wide experimental datasets. Nucleic Acids Res. 47, D607–D613 (2019).

50. Kerrien, S. et al. IntAct--open source resource for molecular interaction data. Nucleic Acids Res. 35, D561–5 (2007).

51. Merali, Z. & Giles, J. Databases in peril. Nature 435, 1010–1011 (2005).

52. O’Brien, K. P., Remm, M. & Sonnhammer, E. L. L. Inparanoid: a comprehensive database of eukaryotic orthologs. Nucleic Acids Res. 33, D476–80 (2005).

53. Altenhoff, A. M. et al. OMA orthology in 2021: website overhaul, conserved isoforms, ancestral gene order and more. Nucleic Acids Res. (2020) doi:10.1093/nar/gkaa1007.

54. Altenhoff, A. M. et al. Standardized benchmarking in the quest for orthologs. Nat. Methods 13, 425–430 (2016).

55. Calderone, A., Castagnoli, L. & Cesareni, G. mentha: a resource for browsing integrated protein-interaction networks. Nat. Methods 10, 690–691 (2013).

56. Mosca, R., Céol, A. & Aloy, P. Interactome3D: adding structural details to protein networks. Nat. Methods 10, 47–53 (2013).

57. Orchard, S. et al. The MIntAct project - IntAct as a common curation platform for 11 molecular interaction databases. Nucleic Acids Res. 42, D358–63 (2014).

58. Oughtred, R. et al. The BioGRID interaction database: 2019 update. Nucleic Acids Res. 47, D529–D541 (2019).

59. Kreimer, A., Borenstein, E., Gophna, U. & Ruppin, E. The evolution of modularity in bacterial metabolic networks. Proc Natl Acad Sci USA 105, 6976–6981 (2008).

60. Nguyen, N. T. T. et al. RSAT 2018: regulatory sequence analysis tools 20th anniversary. Nucleic Acids Res. 46, W209–W214 (2018).

61. Kingsley, R. A. et al. Epidemic multiple drug resistant Salmonella Typhimurium causing invasive disease in sub-Saharan Africa have a distinct genotype. Genome Res. 19, 2279–2287 (2009).

62. Canals, R. et al. Adding function to the genome of African Salmonella Typhimurium ST313 strain D23580. PLoS Biol. 17, e3000059 (2019).

63. Goenawan, I. H., Bryan, K. & Lynn, D. J. DyNet: visualization and analysis of dynamic molecular interaction networks. Bioinformatics 32, 2713–2715 (2016).

64. Alexa, A. & Rahnenfuhrer, J. topGO: Enrichment Analysis for Gene Ontology. https://www.bioconductor.org/packages/release/bioc/html/topGO.html (2021).

65. Wu, T. et al. clusterProfiler 4.0: A universal enrichment tool for interpreting omics data. Innovation (N Y) 2, 100141 (2021).

